# Turbulent-like dynamics in the human brain

**DOI:** 10.1101/865923

**Authors:** Gustavo Deco, Morten L. Kringelbach

## Abstract

Turbulence facilitates fast energy/information transfer across scales in physical systems. These qualities are important for brain function, but it is currently unknown if the dynamic intrinsic backbone of brain also exhibits turbulence. Using large-scale neuroimaging empirical data from 1003 healthy participants, we demonstrate Kuramoto’s amplitude turbulence in human brain dynamics. Furthermore, we build a whole-brain model with coupled oscillators to demonstrate that the best fit to the data corresponds to a region of maximally developed amplitude turbulence, which also corresponds to maximal sensitivity to the processing of external stimulations (information capability). The model shows the economy of anatomy by following the *Exponential Distance Rule* of anatomical connections as a cost-of-wiring principle. This establishes a firm link between turbulence and optimal brain function. Overall, our results reveal a way of analysing and modelling whole-brain dynamics that suggests turbulence as the dynamic intrinsic backbone facilitating large scale network communication.

## Introduction

The study of turbulence remains one of the most exciting unsolved problems of modern physics (Cross and Hohenberg, 1993). Much progress has been made in the field of fluid and oscillator dynamics in terms of understanding and modelling turbulence (Cross and Hohenberg, 1993; Kawamura et al., 2007; Kuramoto, 1984). One of the most relevant aspects of turbulence is the ability to facilitate fast energy transfer across in fluids, the statistical study of which was pioneered by Andrey Kolmogorov (Frisch, 1995; Kolmogorov, 1941a, b). At an abstract level, effective energy transfer can be thought of in terms of efficient information processing. Thus, a key question presents itself, namely are there turbulence-like dynamics in the human brain?

In the context of turbulence in fluid dynamics, Kolmogorov developed his phenomenological theory of turbulence (Kolmogorov, 1941a, b) (see excellent review in (Frisch, 1995)). This introduced the important concept of *structure functions*, based on computing the spatial correlations between any two points in a fluid, which demonstrated and quantified the energy cascades that balance kinetics and viscous dissipation.

Later on, Kuramoto used the theory of coupled oscillators to show turbulence in fluid dynamics (Kuramoto, 1984). Beyond fluid dynamics, coupled oscillators have in general been highly successful for describing large scale brain activity and in particular its metastability ((Cabral et al., 2014; Deco et al., 2017c)). These findings suggest that turbulence could play a role in brain dynamics, and could be important for ensuring efficient information transfer (rather than energy transfer). Specifically, in the coupled oscillator framework, the Kuramoto local order parameter represents a spatial average of the complex phase factor of the local oscillators weighted by the coupling. The level of amplitude turbulence is defined as the standard deviation of the modulus of Kuramoto local order parameter and can be applied to the empirical data of any physical system.

Here, to investigate the presence of turbulence-like traces in human brain dynamics, we combined Kuramoto’s framework for describing turbulence together with Kolmogorov’s concept of structure functions for describing turbulence. We applied this framework to a large Human Connectome Project (HCP) database with neuroimaging data from 1003 healthy human participants. We found that the empirical data shows clear evidence of Kuramoto’s amplitude turbulence (indexed by the local Kuramoto order parameter).

One thing is to observe, however, and another is to truly understand a phenomenon through a causal mechanistic model. The dynamics of the human brain has been described using a plethora of whole-brain models, which include biophysical realistic models (Chaudhuri et al., 2015; Deco and Jirsa, 2012; Demirtas et al., 2019; Demirtas et al., 2017; Ghosh et al., 2008; Honey et al., 2009; Izhikevich and Edelman, 2008) and models of coupled oscillators (Cabral et al., 2014; Deco et al., 2017b; Deco et al., 2017c). Yet, no one has yet investigated whether these dynamics show traces of turbulence.

Therefore we used a whole-brain model utilising Stuart-Landau (also known as Hopf) oscillators (Deco et al., 2017c) and using the *Exponential Distance Rule* of anatomical connections as a cost-of-wiring principle. This demonstrated the economy of anatomy and showed that, at the dynamical working point of the whole-brain model optimally fitting the empirical data, the system shows not just Kuramoto’s amplitude turbulence but *maximal* amplitude turbulence. Even further, we generalised the concept of susceptibility which measures the sensitivity of the brain to the processing of external stimulation, to define a measure of the information capability of the whole-brain model. The information capability is designed to capture how different external stimulations of the model are encoded in the elicited dynamics (see Methods). Remarkably, at the dynamical working point of the model fitting the data where there is maximal amplitude turbulence, we also found maximal information capability.

This framework also allowed us to investigate the differences between the amplitude turbulence found in resting state and in the seven behavioural tasks found in HCP dataset. The results show that they share a turbulent core but that the long-distance correlations show task-specific increases in higher-order brain regions outside the turbulent core.

Finally, given that we have shown that turbulence is the dynamic intrinsic backbone facilitating large scale network communication, we also investigated if there are power laws in empirical brain dynamics similar to those found by Kolmogorov in the structure functions of fluid dynamics. In our case, however, such power laws would be evidence of the presence of a cascade of efficient information processing across scales. We found power laws in the turbulent core, tentatively named the ‘inertial subrange’ similar to those found in fluid dynamics, and which similarly appear to be homogeneous isotropic, i.e. with average properties that are both independent of position and direction.

## Results

In order to demonstrate turbulence in human brain dynamics we combined the seminal insights and methods of Kolmogorov (Kolmogorov, 1941a, b) and Kuramoto (1984). The study of turbulence in fluid dynamics (see left panel of **Figure 1A**) was strongly influenced by Richardson’s concept of cascaded eddies reflecting the energy transfer (see cartoon in right panel of **Figure 1A**), where the hierarchical organisation of different sizes of eddies is schematised for the turbulent so-called ‘inertial subrange’, ie the range where turbulence kinetic energy is transferred from larger to smaller scales without loss (see shaded areas in **Figures 1A** and **1B)**. Subsequently, this inspired Kolmogorov to create his phenomenological theory of turbulence based on the concept of structure functions. For fluid dynamics, he demonstrated the existence of power laws in the inertial subrange where the structure functions show a universal scaling of the spatial scale, r, given by 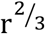 (left panel of **Figure 1B**) and an energy scaling of k, the associated wave number of the spectral scale given by 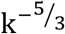 (right panel of **Figure 1B**).

**Figure 1.**
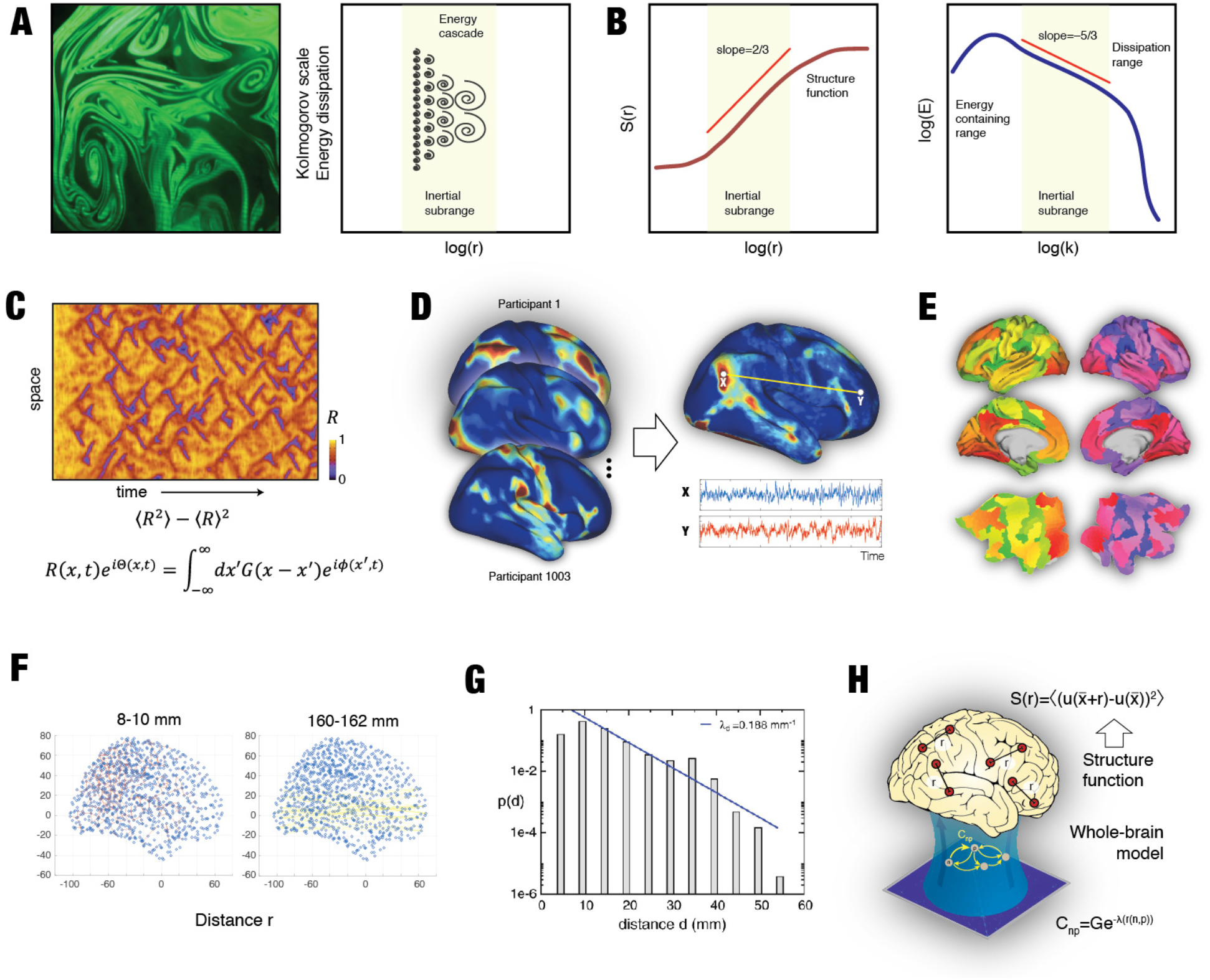
Measuring turbulence in fluid dynamics and in human brain activity. **A)** The study of turbulence in fluid dynamics was pioneered by Kolmogorov’s phenomenological theory of turbulence which is based on the concept of structure functions. In turn, this was inspired by Richardson’s concept of cascaded eddies. The left panel shows a snapshot of turbulence in a real physical system with different sizes of eddies, whose hierarchical organisation is schematised for the inertial subrange in the right panel. **B)** In fluid dynamics, as shown in the cartoon, power laws are found in an inertial subrange where the structure functions show a universal scaling of 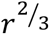 (left panel) and an energy scaling of 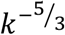 (right panel), where r is the spatial scale and k the associated wave number of the spectral scale. This power law behaviour reflects the energy transfer cascade found in turbulence. **C)** Fluid dynamics can equally well be modelled by coupled oscillators as shown by Kuramoto (1984). He defined a local order parameter, representing a spatial average of the complex phase factor of the local oscillators weighted by the coupling. The standard deviation of the modulus of this measure defines the level of amplitude turbulence, which is shown in the adapted figure for a ring of Stuart-Landau oscillator system (Kawamura et al., 2007). This concept is not only valid for coupled oscillator modelling but also can be used to detect turbulence in a given system in a model-free way. **D)** Here, to detect the presence of amplitude turbulence in human brain activity, we used state-of-the-art resting state data from a large set of 1003 healthy human participants in the Human Connectome Project (HCP) database. **E)** We extracted the timeseries from each the 1000 parcels in the fine-grained Schaefer parcellation, here shown as slices in MNI space and on the surface of the HCP CIFTI space. **F)** The function structure is based on the functional correlations between pairs with equal Euclidean distance, r, in MNI space. Here we show two examples of the pairs with r=8-10 mm (top) and r=160-162 mm (bottom). **G)** It has been that most of the underlying brain connectivity follows the exponential decay described by the Exponential Distance Rule (Ercsey-Ravasz et al., 2013). The figure shows the histogram of interareal projection length for all labeled neurons (n = 6,494,974) in a massive tract tracing study in non-human primates. The blue line shows the exponential fit with a decay rate 0.188mm^-1^. **H)** We used this anatomical basis in a whole-brain model based on Stuart-Landau oscillators (Deco et al., 2017c) aiming to establish the causal mechanisms underlying the emergence of turbulence.

Another way to describe turbulence in fluid dynamics was proposed by Kuramoto (1984), who defined a local order parameter, representing a spatial average of the complex phase factor of the local oscillators weighted by the coupling. The amplitude turbulence is simply given by the standard deviation of the modulus of this measure. An example of this is shown in **Figure 1C** for a ring of Stuart-Landau oscillator system (Kawamura et al., 2007).

We used the state-of-the-art resting state data from a large set of 1003 healthy human participants in the Human Connectome Project (HCP) database (see **Figure 1D** and Methods), extracting the timeseries from each the 1000 parcels in the fine-grained Schaefer parcellation (Schaefer et al., 2018) (**Figure 1E**). The empirical data was minimally pre-processed according to the HCP protocol and subsequently filtered in the narrow relevant band between 0.008 and 0.08 Hz, detrended and z-scored (see Methods). We computed the function structure as the functional correlations between pairs with equal Euclidean distance, r, in MNI space (**Figure 1F**). We combined Kolmogorov’s structure functions with Kuramoto’s local order parameter to demonstrate amplitude turbulence. Finally, we created a whole-brain model using simplified brain connectivity following the Exponential Distance Rule (Ercsey-Ravasz et al., 2013; Markov et al., 2013; Markov et al., 2014) based on massive tract tracing studies in non-human primates (**Figure 1G**). This whole-brain model was based on Stuart-Landau (also called Hopf) oscillators (Deco et al., 2017c) aiming to establish the causal mechanisms underlying the emergence of turbulence (**Figure 1H**).

### Amplitude turbulence in empirical brain dynamics

We computed the local Kuramoto order parameter, R, for the empirical brain resting data of 1200 data points from the 1003 HCP participants and compared this to surrogate data, a shuffled version maintaining the spatiotemporal characteristics of the empirical data (Kantz and Schreiber, 1997). Amplitude turbulence, D, is defined as the standard deviation of the modulus of the local Kuramoto order parameter *R* across time and space (Kawamura et al., 2007). **Figure 2A** (left panel) shows a box plot of the statistical significant difference (p<0.001, two-sided Wilcoxon rank sum test) between empirical and surrogate data. Furthermore, to ascertain the absence of regular spatiotemporal patterns in the empirical data, we computed the autocorrelation of the local Kuramoto order parameter, R, across space and time (middle and right panel, respectively), which show a rapid decay as expected in turbulence.

**Figure 2.**
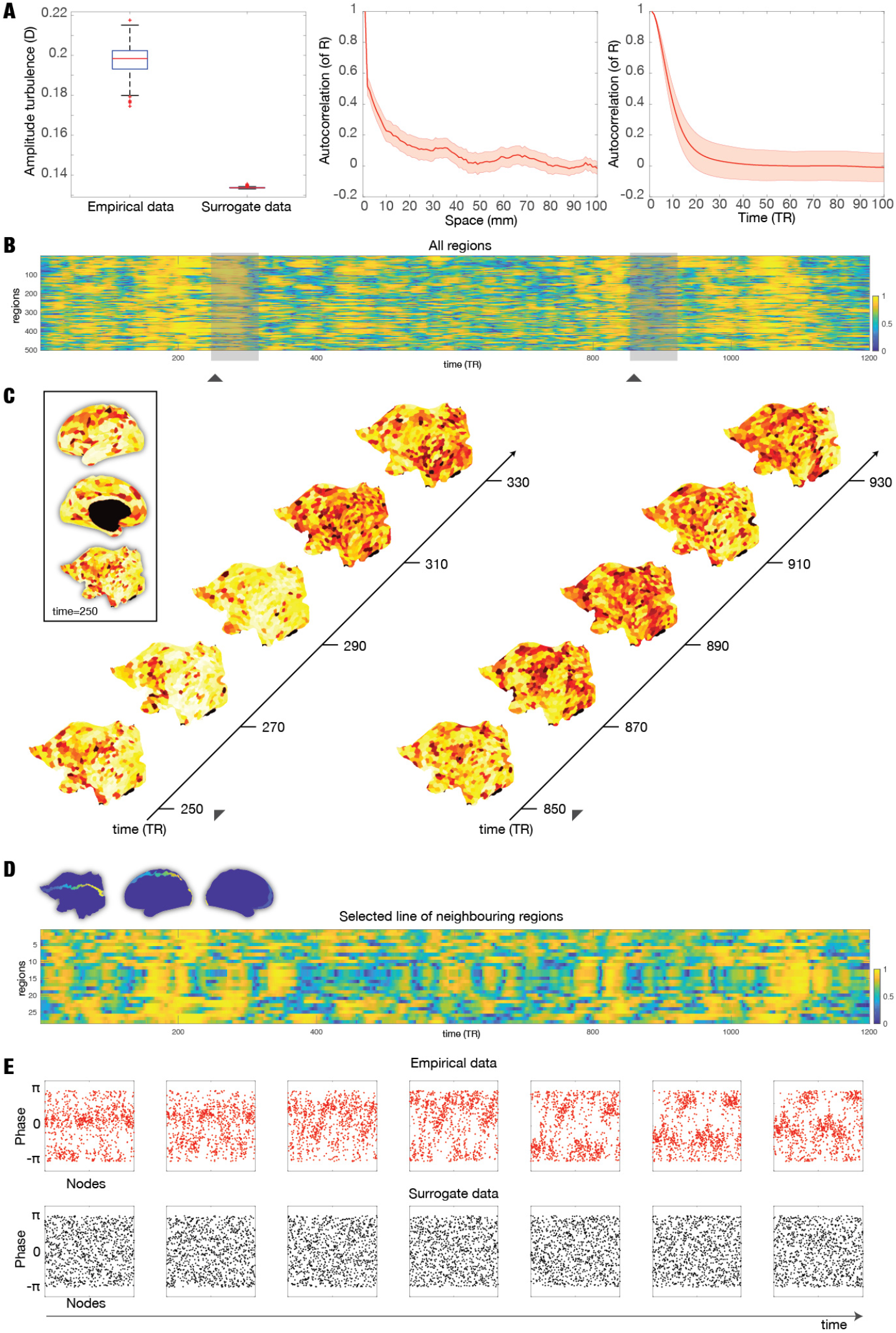
Amplitude turbulence in empirical data. **A)** The left panel shows a boxplot of the amplitude turbulence, D, computed on the empirical resting state data of the 1003 HCP participants and on the carefully matched surrogate data. These are significantly different (P<0.001, two-sided Wilcoxon rank sum test). The middle and right panel shows the autocorrelation of the local Kuramoto order parameter, R, across space and time, respectively. The rapid decay demonstrates absence of regular spatiotemporal patterns in the empirical data. **B)** The figure visualises the change over time and space of the local Kuramoto order parameter, R, reflecting amplitude turbulence in a single participant. Amplitude turbulence can be clearly seen in the 2D plot of all 500 parcels in the left hemisphere over the 1200 timepoints. **C)** This can be appreciated from the continuous snapshots for two segments separated in time (left and right parts) rendered on a flatmap of the hemisphere (see insert with renderings of a single snapshot on the inflated and flatmapped cortex). Furthermore, the full spatiotemporal evolution can be appreciated in the video (found in the supplementary material, **Video S1**) over the full 1200 timepoints of the full resting state session. **D)** The synchronisation of clusters over time is dependent on the neighbourhood and so to further visualise the spatiotemporal evolution of amplitude turbulence, we show a 2D plot of 26 neighbouring parcels running from the front to the back of the brain (see blue insert). **E)** The figures further demonstrate the presence of turbulence by plotting consecutive snapshots over time of the phases of all brain regions for both the empirical data (top) and the surrogate data (bottom). This clearly shows the absence of structure in the surrogate data and clustering resembling vortices in the empirical data (although note that the regions are simply ordered in their original space similar to B, and therefore potentially show less of the neighbourhood effect shown in D).

It is instructive to visualise the change over time and space of the local Kuramoto order parameter, R. **Figure 2B** shows the spatiotemporal evolution of amplitude turbulence in empirical data of a single participant in a 2D plot of all 500 parcels in the left hemisphere over the 1200 timepoints. Given that this is a 1D representation of a 3D space, the 500 parcels are not ordered in terms of spatial neighbourhood and therefore this does not represent the true spatiotemporal evolution of amplitude turbulence. Instead to appreciate the synchronisation of neighbouring clusters over time, **Figure 2C** shows snapshots for two segments separated in time (the left and right parts marked on the 2D plot) rendered on a flatmap of the hemisphere. The evolution of turbulence in the empirical data across space and time is even clearer in **Video S1** in the supplementary material, which shows the full spatiotemporal evolution over the full 1200 timepoints of the full resting state session. Remarkably, the evolution of amplitude turbulence in terms of R, where spatial neighbourhood is conserved, closely resembles the typical turbulence found in fluid dynamics and oscillators, which can now be directly compared with the theoretical ring results of Kawamura and colleagues (Kawamura et al., 2007). Furthermore, **Figure 2D** shows this in another way by plotting only 26 neighbouring parcels running from the front to the back of the brain.

Finally, similar to Kawamura and colleagues (Kawamura et al., 2007), **Figure 2E** plots consecutive snapshots over time of the phases of all brain regions for both the empirical data (top) and the surrogate data (bottom). This figure convincingly demonstrates the absence of structure in the surrogate data and clustering resembling vortices in the empirical data.

### Mechanistic origins of turbulent human brain dynamics

We wanted to understand the causal mechanistic principles underlying the emergence of turbulence in brain dynamics. Turbulence has been described by Kuramoto using Stuart-Landau oscillators (Kuramoto, 1984), and the very same oscillators have successfully been used in whole-brain models modelling human brain activity, although these models are usually named after the Hopf bifurcation, given the fact that the Stuart-Landau oscillator expressed mathematically is the normal form of the Hopf bifurcation (Deco et al., 2017c). Therefore Hopf whole-brain models are highly suitable for elucidating the underlying mechanisms of turbulence in brain dynamics.

Whole-brain modelling couples local dynamics between different brain regions through their anatomical structural connectivity, which usually obtained from tractography estimated with dMRI. On the other hand, massive tract tracing studies in non-human primates have shown that the core of anatomical structural brain connectivity can be fairly well described by a simple rule, the Exponential Distance Rule, given by C_np_ = e^−λ(r(n,p))^ (Ercsey-Ravasz et al., 2013) (see **Figure 1G** and Methods). Here C_np_ is the anatomical coupling between brain region n and p, and λ is the exponential decay of the connectivity as a function of the distance, i.e. r(n, p), which is simply the Euclidean distance between brain regions in MNI space.

**Figure 3A** shows the close relationship between the empirical HCP dMRI tractography of the human brain and the exponential distance rule. Specifically, the figure shows a plot of the fibre densities between the pairs of regions in the Schaefer parcellations as a function of the Euclidian distance, r, between the nodes. The blue line represents dMRI tractography and the red line represents the fitted exponential distance rule. The subpanels show the structural connectivity matrices for the empirical dMRI tractography (left) and the fitted exponential distance rule (right), at the optimal λ=0.18 mm^−1^ when fitting the dMRI connectivity data to the underlying exponential function. These matrices are remarkably similar reflecting the excellent level of fitting.

**Figure 3.**
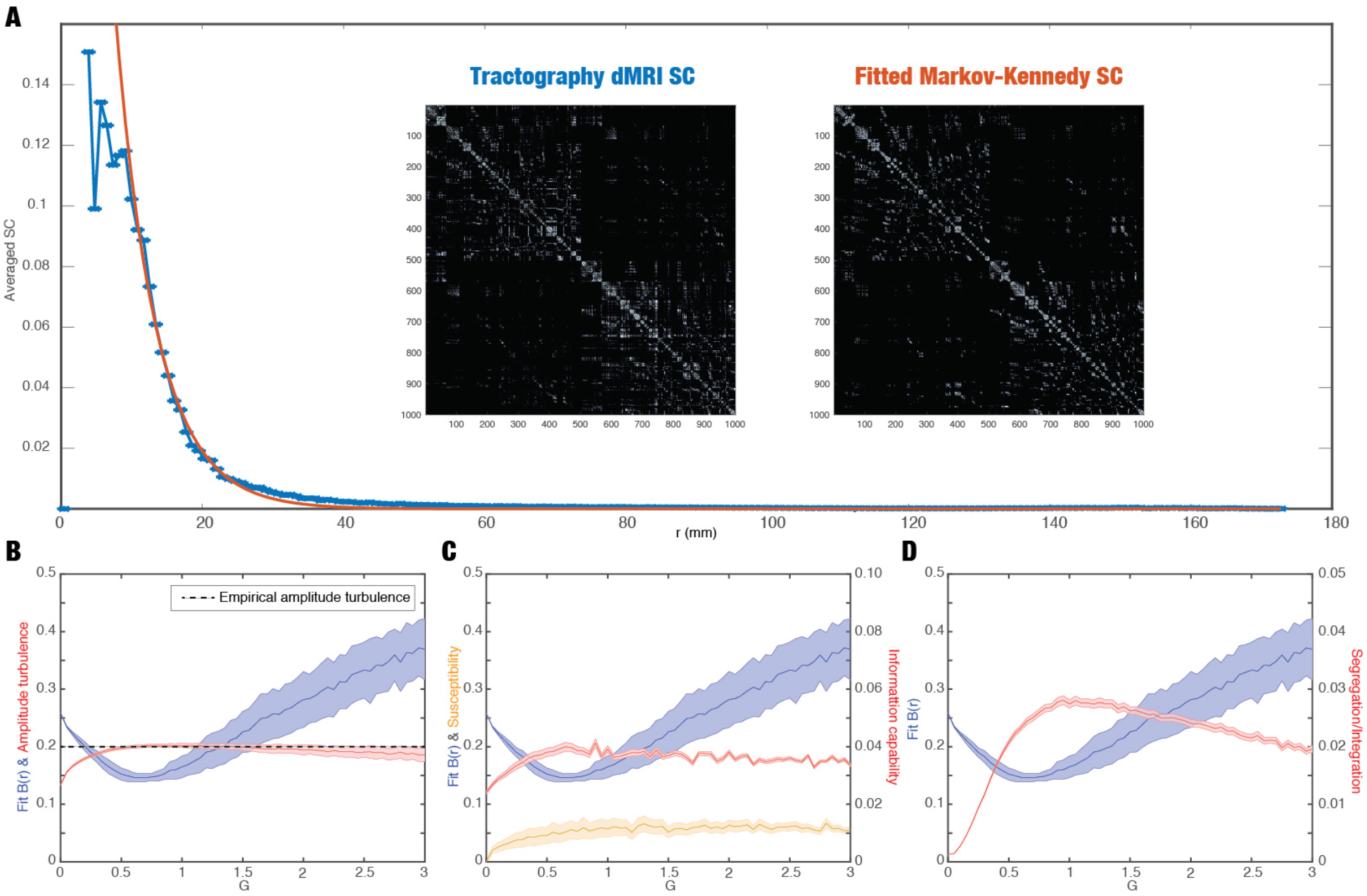
Whole-brain modelling demonstrating turbulence in the empirical data. **A)** The Exponential Distance Rule is also evident in the empirical HCP dMRI tractography of the human brain, as shown by the fibre densities between the pairs of regions in the Schaefer parcellations as a function of the Euclidian distance between the nodes with blue showing dMRI tractography and red line showing the fitted Exponential Distance Rule at the optimal λ=0.18 mm^−1^. The remarkable similarity can be appreciated by comparing the two subpanels. On the left is shown the structural connectivity matrices for the empirical dMRI tractography and on the right the optimally fitted Exponential Distance Rule connectivity, which was used as the basis for the whole-brain model. **B)** The figure shows the whole brain fit of the root squared error between the empirical and simulated B(r) in the inertial subrange as a function of the global coupling parameter G (black). The model shows amplitude turbulence (red line, defined in Methods) in a broad range of G but maximal amplitude turbulence is found at the optimal working point fitting the data (G=0.8). The dotted line shows the amplitude turbulence estimated from the empirical data, and it is interesting that the model at the optimal working point also corresponds to this value. **C)** The maximal amplitude turbulence is likely to reflect an optimal level of information processing, which we quantify in a measure of information capability, a meaningful extension of the standard concept of susceptibility (see Methods). As can be seen the maximum of information capability (red line) is found at G=0.8 which corresponds to the optimal fitting of the whole-brain model to the empirical data (black line) and maximal amplitude turbulence. In contrast, the simple measure of susceptibility (orange line) is high but not maximal at the working point. **D)** Interestingly at the optimal point where the whole-brain model fits the empirical data (black line) and shows maximal amplitude turbulence and information capability, we also find an optimal balance between segregation/integration (red line) as a function of G.

This fact simplifies the fitting of a Hopf whole-brain model to the empirical functional data. We built a Hopf whole-brain model using the exponential distance rule (with the empirically derived λ=0.18 mm^−1^) (**Figure 1H**). As the fitting function, we used Kolmogorov’s concept of *structure functions* of a variable *u* (in turbulence usually a transversal or longitudinal velocity), which is defined here (**Figure 1H** and in the Methods) as 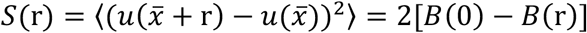, where the basic spatial correlations of two points separated by an Euclidean distance *r*, are given by 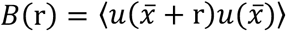). Here, we use variable *u* to denote the spatiotemporal fMRI BOLD signals from our analysis of the whole-brain dynamics of the HCP resting state data. The symbol 〈 〉 refers to the average across the spatial location 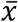 of the nodes and time. Please note that many pairs of nodes across the brain have the same Euclidean distance, *r*, and are thus averaged, as indicated by the notation 〈 〉. As can be seen from the two equations, *S*(*r*) and *B*(*r*), they are basically characterising the evolution of the functional connectivity as function of the Euclidean distance between equally distant nodes. This is different from the usual definition of functional connectivity which is not measured as a function of the distance. Thus, we studied the whole brain fit of the root squared error between the empirical and simulated *B*(*r*) as a function of the global coupling parameter, G (see Methods). This global coupling parameter regulates the effectivity of the interactions between regions.

**Figure 3B** shows the evolution of the amplitude turbulence, D, (red line) and the level of model fitting (blue line) as a function of the global coupling parameter, G (see Methods). We find that the amplitude turbulence, D, is increasing with G and reach a plateau until tapering off at high levels. This means that the model exhibit turbulence at a broad range of global coupling strengths. Remarkably, however, the maximum of amplitude turbulence, D, is found at the optimal working point at G = 0.8, where the model fits the empirical data; specifically B(r), the spatial correlation function of two nodes. The fact that we get a maximum of turbulence at the working point could suggest that the level of turbulence is reflecting the information capabilities of the brain. Furthermore, the level of amplitude turbulence for the empirical data is plotted by a dotted line, which corresponds to the maximum. In other words, at the optimal working point, the model is not only reaching its maximum but this is also corresponding the empirical value. As a note, **Figure S1** shows that using the traditional strategy of fitting using the correlation between empirical and simulated functional connectivity matrices (red line) is not informative for constraining the model. This shows the usefulness of defining functional connectivity as a function of equally distanced nodes as is defined in the structure function definitions of *S*(r) and *B*(r).

In order to investigate the role of turbulence on fitting the model, we were inspired by the findings by Kawamura, Nakao and Kuramoto (Kawamura et al., 2007), who manipulated the shear parameter (see Methods). **Figure S2** shows the results of systematically changing the shear parameter, β, and the resulting changes in amplitude turbulence (red line) and the level of fitting (black line). Increasing the shear parameter leads to a worsening of the level of fitting, which is the error of the estimation of B(r). Nevertheless, the shear parameter is strongly affecting the amplitude turbulence, and it is interesting to observe that when this decreases so the level of fitting get worse, establishing the relevance of turbulence for fitting the model.

We investigated the possibility that the level of turbulence reflects *information capability* of the brain by generalising the concept of *susceptibility*, which is defined as the sensitivity of the whole-brain model to the processing of external stimulations (see Methods). On the other hand, the *information capability* of the whole-brain model is defined as the standard deviation across trials of the difference between the perturbed and unperturbed mean of the modulus of the local order parameter across time, averaged over all brain nodes (see Methods). This is easy to implement in the Hopf whole-brain model (Deco et al., 2017a; Deco et al., 2019), where perturbations can be introduced by changing the local bifurcation parameter a_n_ of each brain node n (see Methods). For each value of G, we perturbed the whole-brain model 200 times with random parameters for the local bifurcation parameter a_n_.

**Figure 3C** shows that the maximum of information capability (red line) is found at G=0.9 which corresponds to the optimal fitting of the whole-brain model to the empirical data (blue line). This clearly demonstrates that maximal turbulence is directly associated with information capability at the working point of the whole-brain model fitting the empirical data and thus presumably reflecting optimal information processing. In contrast, as is shown by the pink line, the simple measure of susceptibility is not maximal at this working point (but high) and in fact does not show a maximum in the range shown for G.

Further probing the question of optimal information processing, we measured the integration and segregation of the whole-brain model in the whole range of global coupling (as before, see Methods). Integration is measured as the mean functional correlation and segregation is measured by the level of modularity of the functional connectivity (see Methods for precise definition). **Figure 3D** shows the combined measure of segregation/integration (red line) G. As can be seen the maximum is around the optimal fitting of the whole-brain model to the empirical data (blue line). This suggests the normal brain is also at its highest level of being able to segregate and integrate information (Deco et al., 2015).

### Spatiotemporal evolution of amplitude turbulence in whole-brain model dynamics

Deeper insights into the spatiotemporal evolution of amplitude turbulence can be gained from studying the Hopf whole-brain model for different coupling strengths showing different levels of amplitude turbulence. **Figure 4A** shows 2D plots of the spatiotemporal evolution of the local Kuramoto order parameter, R, reflecting different levels of turbulence in the model for four different coupling strengths (G=0, G=0.4, G=0.8, and G=3.0). Similar to **Figure 2**, we plot the level of R for all 500 parcels in the left hemisphere over 1200 timepoints. The optimal working point (G=0.8) is highlighted and shows maximal turbulence as can be appreciated by comparing the level of variability of R in the other three 2D plots. Please also note how the uncoupled case (G=0) resembles random spatiotemporal dynamics, while the two cases demonstrate various degrees of turbulence (as shown by the values of D, the standard deviation of R, in **Figure 3B**). **Figure 4B** shows the corresponding 2D plots of the spatiotemporal evolution of R in 26 neighbouring parcels running from the front to the back of the brain (similar to the plot in **Figure 2D**).

**Figure 4.**
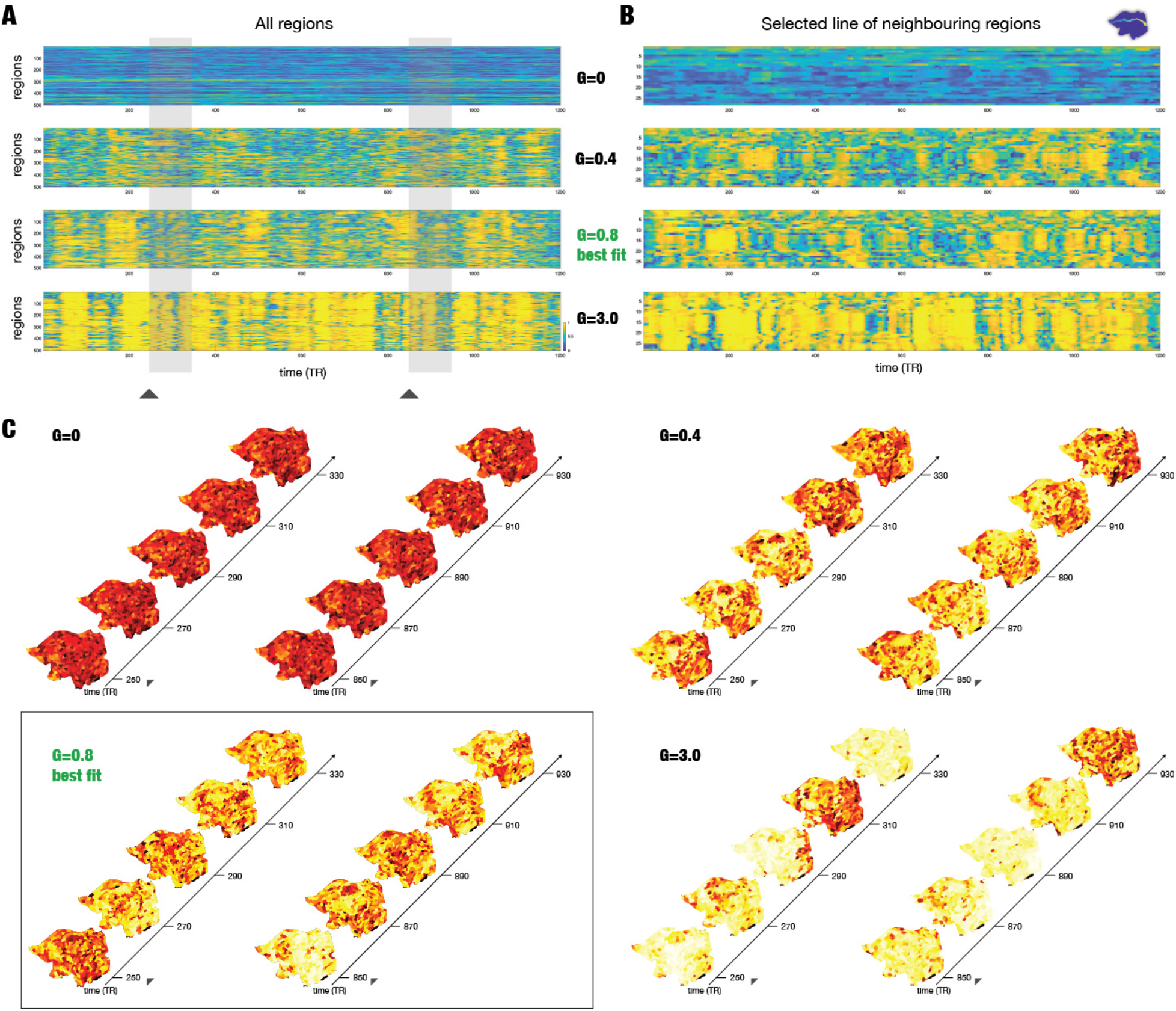
The spatiotemporal evolution of whole-brain model for different coupling strengths showing different levels of amplitude turbulence. **A)** For four different coupling strengths (G=0, G=0.4, G=0.8, and G=3.0), we show a 2D plot of the spatiotemporal evolution of the local Kuramoto order parameter, R, reflecting different levels of turbulence in the model (for all 500 parcels in the left hemisphere over 1200 timepoints). The highlighted optimal working point (G=0.8 in red) is showing maximal turbulence as can be appreciated by comparing to the other three 2D plots. **B)** Similar to Figure 4C, for all four values of coupling strengths, G, we show 2D plots of the spatiotemporal evolution of amplitude turbulence in 26 neighbouring parcels running from the front to the back of the brain. **C)** Similarly, we show continuous snapshots for two segments of the model at G=0, G=0.4, G=0.8, and G=3.0, separated in time (left and right parts) rendered on a flatmap of the hemisphere. Furthermore, the full spatiotemporal evolution of each can be appreciated in the videos for each G (found in the supplementary material, **Videos S2-S5**) over the full 1200 timepoints of the full resting state session.

The supplementary material contains four videos (**Videos S2-S5**) of the full spatiotemporal evolution of amplitude turbulence in one hemisphere across the 1200 timepoints of the full resting state session for each G. For the optimal working point fitting the empirical data (G=0.8), **Figure 4C** shows snapshots of for two segments of separated in time rendered on a flatmap of the hemisphere for each the four values of G. Again, it is remarkable how the spatiotemporal patterns generated by the wholebrain model at for the optimal working point fitting the data (G=0.8) resemble the amplitude turbulence found in the empirical brain activity. It is also clear that there is amplitude turbulence for other values of G, but as shown in **Figure 3B**, the maximal value of turbulence is observed at the working point of the model fitting the empirical data.

### Differences between task and resting

These results demonstrate amplitude turbulence in the brain dynamics of resting state. We were interested in investigating how turbulence is controlled when performing different cognitive tasks. To address this question, we studied the seven HCP tasks and contrasted these results with rest.

First, we established a spatial map of the most significant correlations in resting state in order to have this as a reference for the analysis of the tasks. **Figure 5A** shows the group average functional connectivity (FC) correlation matrix for all 1000 parcels across all 1003 participants in the resting state. We calculated the global brain connectivity (GBC) of the FC matrix by calculating the average functional connectivity of each region with all other regions (the mean value of each row across the columns). Right panel of **Figure 5B** shows a rendering of GBC thresholded at the 80% quantile.

**Figure 5.**
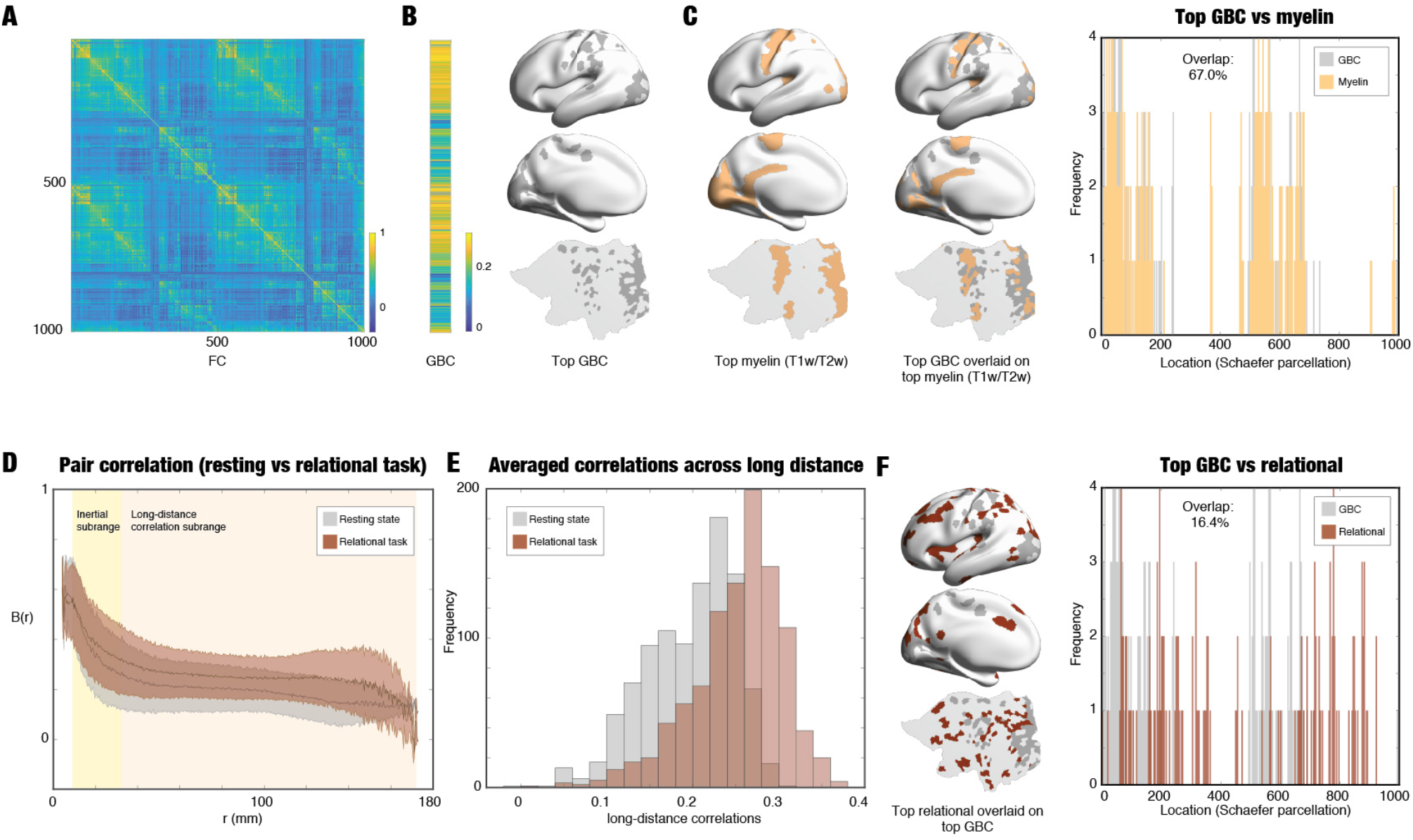
Discovering the functional exceptions driving the turbulent core in cognitive tasks. **A)** The figure shows the functional connectivity in resting state between the 1000 regions in the Schaefer1000 parcellation averaged across all HCP 1003 participants. **B)** Similarly, from this we computed the global brain connectivity (GBC) as the mean correlation between each region with the rest of the brain (Demirtas et al., 2019) which characterises the node-level connectivity. The left panel shows the GBC vector (averaged over all participants) while the right panel shows a rendering on the brain of position of the top 20% quantile GBC regions. The top part shows them rendered on the left hemisphere, the middle part the midline while the bottom part shows a rendering on a flat map of the left hemisphere. **C)** The left panel shows the top regions with myelination (T1w/T2w) rendered on the brain. As can be seen in the middle panel, there is a strong spatial overlap between the top GBC regions and the top regions with myelination. This can also be seen in the right panel of the overlapping histograms of the top GBC and top myelin regions as indexed by the spatial location (Schaefer parcellation number), which shows a 46.5% overlap. **D)** Example of the pipeline finding the functional exceptions applied to HCP relational task (see Methods). Here is shown the contrast between the relational task (brown) and the resting state (grey), with the shaded error showing the dispersion across nodes, i.e. all pairs across all participants. The inertial subrange (r=[8.13 33.82] mm) is highlighted with a light yellow background, while the long-distance correlation subrange (r>33.82 mm) is shown on a light grey background. As can been clearly seen, the long-range correlations are mainly increased in task, whereas the inertial subrange correlations remain unchanged (p<0.001, Wilcoxon rank sum). **E)** We show a histogram of the difference of the average correlation for each spatial location across the long-distance subrange. The histogram for task (in brown) is clearly showing higher correlations than the histogram for resting (in grey, p<0.001, Wilcoxon rank sum). F) We found the most changed long-distance regions in the relational task by thresholding the pair correlation by the maximum pair correlation of the resting condition. The left panel shows a rendering of the top changing regions in the relational task overlaid on the top GBC regions. The overlap is very low (18.2%) as can be seen in the right panel, which shows the overlapping histograms of the top relational (red) and top GBC (blue) regions as indexed by the spatial location. This is strong evidence that the most changed regions in task are complementary to the unchanged resting GBC regions.

Remarkably, as shown in **Figure 5C**, there is a strong spatial overlaps with the myelination measured with T1w/T2w (Glasser et al., 2014; Glasser and Van Essen, 2011) thresholded at a similar 80% quantile (left subpanel). This can be seen in the middle panel of **Figure 5C**, where GBC is overlaid on spatial myelin map, and in the right panel of the overlapping histograms of the top GBC and top myelin regions as indexed by the spatial location (indexed by the Schaefer parcellation number), which are showing a 67.0% overlap. This means that the backbone of resting state processing builds a functional core in primarily visual, auditory and somatomotor regions.

Second, for rest and each task we compare the correlation function *B*(*r*) as a function of the distance *r* (see Methods). As a representative example, **Figure 5D** shows the contrast between the HCP relational task (red, see Methods) and the resting state (grey), with the shaded error showing the dispersion across nodes, i.e. all pairs across all participants. As can be seen from **Figure 5D**, it would appear that the task and rest are similar in a subrange of values for *r* and that precisely in this range, there is a power law. We will return to this power law behaviour in the next section. Inspired by Kolmogorov, we use the term *‘inertial subrange’* to refer to the range *r*=[8.13 33.82] mm (light yellow background), which is the functional core, where task and rest are similar. In contrast, we use the term *‘long-distance correlation subrange’* for r>33.82 mm (light grey background), which is outside the functional core and where task and rest are dissimilar. As can be seen, the inertial subrange is mainly unaffected, but the long-distance correlations are significantly increased in the relational task (p<0.001, Wilcoxon rank sum, and for all other tasks, not shown).

**Figure 5E** shows the histograms of rest (grey) and relational task (red) of correlations averaged across the long-distance subrange. As can be seen clearly, the two distributions are significantly different (p<0.001, Wilcoxon rank sum) and there is a group of task-specific regions that show larger correlations than the maximum of resting state (which is equally true for the six other HCP tasks). **Figure 5F** (left subpanel) shows the spatial maps of the relational task-specific regions are found in higher-order brain regions (in red) outside the functional core, and overlaid on the maps for the thresholded GBC map from resting state (in grey). Remarkably, the overlap is very low (16.4%) as can be seen in the right panel, which shows the overlapping histograms of the top relational (red) and top GBC (grey) regions as indexed by the spatial location. This finding demonstrates that the task-specific regions are taken from the long-range correlations that serve to control the unaffected functional core.

**Figure 6A** shows the same procedure of thresholding the correlations averaged over the long-distance subrange (for the maximum value of the resting state long-distance correlations) but now applied to all seven HCP tasks (relational, red; gambling, green; emotion, light blue; working memory (wm), light red; social, pink; language, blue and motor, purple). These are rendered to visualise the task-specific regions for each task, overlaid on the thresholded GBC map from resting state (grey regions). The regions are found in higher-order regions of the frontal, orbitofrontal, parietal, temporal, insular and midline frontal and cingulate cortices.

Importantly, for all tasks there are hardly any overlaps with the resting GBC maps strongly suggestive of significant role of long-distance connections in controlling the unaffected common turbulent core. Interesting, as expected, each of the tasks use different, yet overlapping higher-order regions to perform the relevant cognitive task.

**Figure 6.**
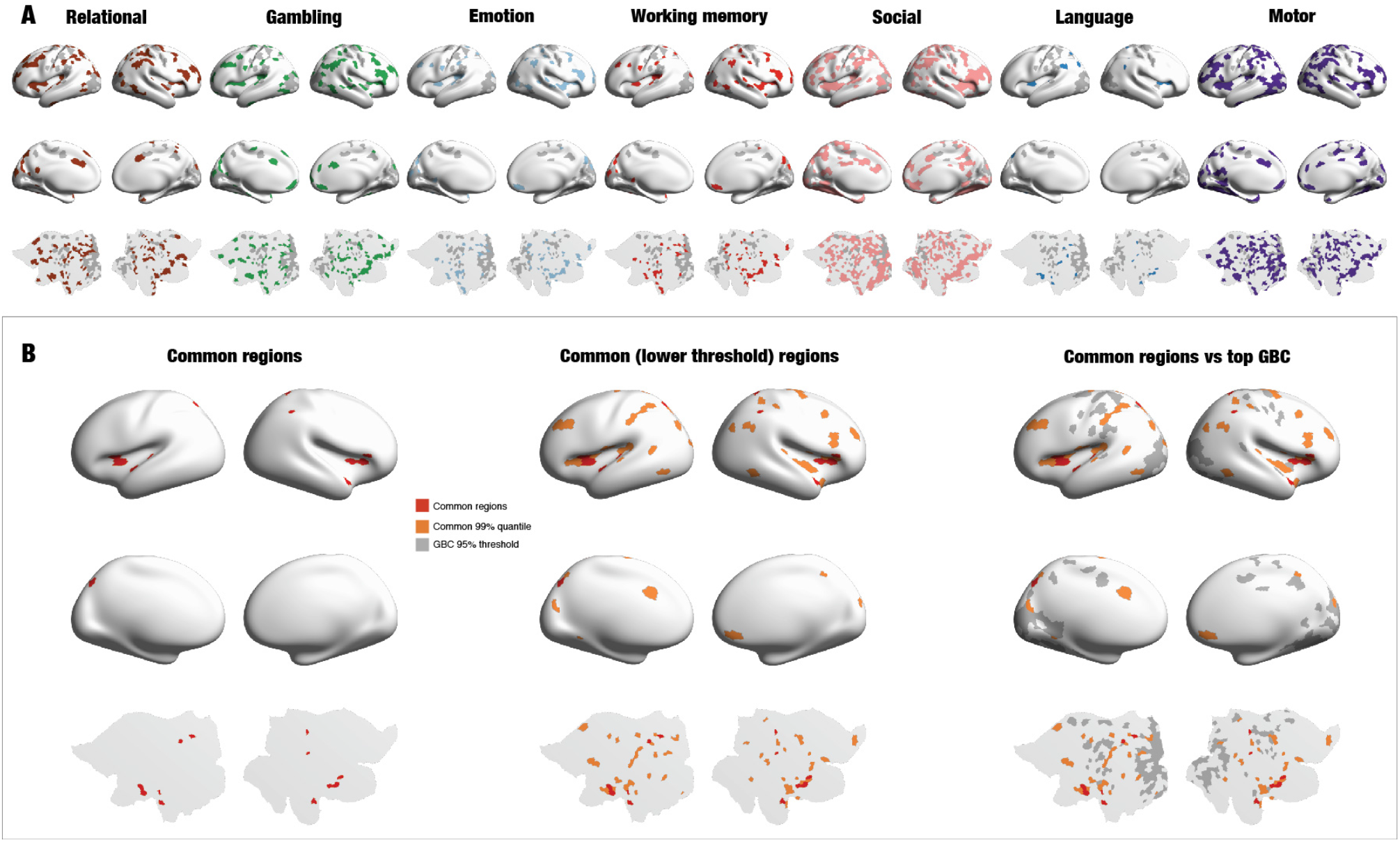
Comparing task-specific exceptions across seven tasks. **A)** The same pipeline shown in Figure 6 was applied to all seven HCP tasks (relational, gambling, emotion, working memory (wm), social, language and motor). These are rendered on side views, midline views and flat map of the left hemisphere to visualise the task-specific regions for each task, overlaid on the thresholded GBC map from the resting state (dark blue). **B)** The overlap of task-specific exceptions is quantified by computing the intersection between task-specific regions by thresholding of the seven tasks at two thresholds: max (leftmost panel, red) and 99% quantile (middle panel, orange) of the resting state long-distance correlations). These are then overlaid on the GBC map (right panel, grey). This overlap could correspond to a “cognitive backbone” which is needed to control the turbulent core processing.

**Figure 6B** shows a quantification of the overlap of task-specific exceptions by computing the intersection between task-specific regions by thresholding of the seven tasks at two thresholds: max (leftmost panel, red) and 99% quantile (middle panel, orange) of the resting state long-distance correlations). These are then overlaid on the GBC map (right panel, grey). The common regions are compatible with the large literature on higher order functional regions. The common regions are known to engage a network of brain regions in the ventromedial prefrontal/orbitofrontal, insular, mid-cingulate and dorsolateral prefrontal cortices. We propose that this overlap could correspond to a “cognitive control network” which is needed to control the turbulence in the functional core processing.

### Exploring the functional core and power law in the empirical data

The important result that the functional core is the underlying backbone for information processing leaves open the important question of whether this shows a power law similar to that found in fluid dynamics, which would be indicative of an information cascade. The existence of such a power law does not, of course, demonstrate the existence of turbulence but provides consistent evidence in support of our main findings of turbulence in the human brain demonstrated using Kuramoto’s oscillator framework in the empirical data and in the Hopf whole-brain models of the data. Other studies have shown power laws in human brain data in the context of criticality which could be consistent with turbulence but is not definite proof (Cocchi et al., 2017; Shew and Plenz, 2013).

We explored whether power laws exist in the inertial subrange sustaining the functional core for both *S*(*r*) and *B*(*r*). We plot both in log-log plots and fit a straight line in the relevant range of r. The slope of the straight line is the exponent of the power law but note that shifting of functions *S* and *B*. Averaging across participants, **Figure 7A** shows the structure function *S*(*r*), plotted in a log-log plot.

We clearly observe an *inertial subrange*, inspired by the similar observation in fluid dynamics, where a power law is found (see Kolmogorov’s law in **Figure 1B**). Our results show a power scaling law with an exponent of approximately 1/2 in the range between *r*=8.13 mm and *r*=33.82 mm in the inertial subrange coinciding with the functional core (shown in the shaded areas in **Figures 7A-C**).

**Figure 7.**
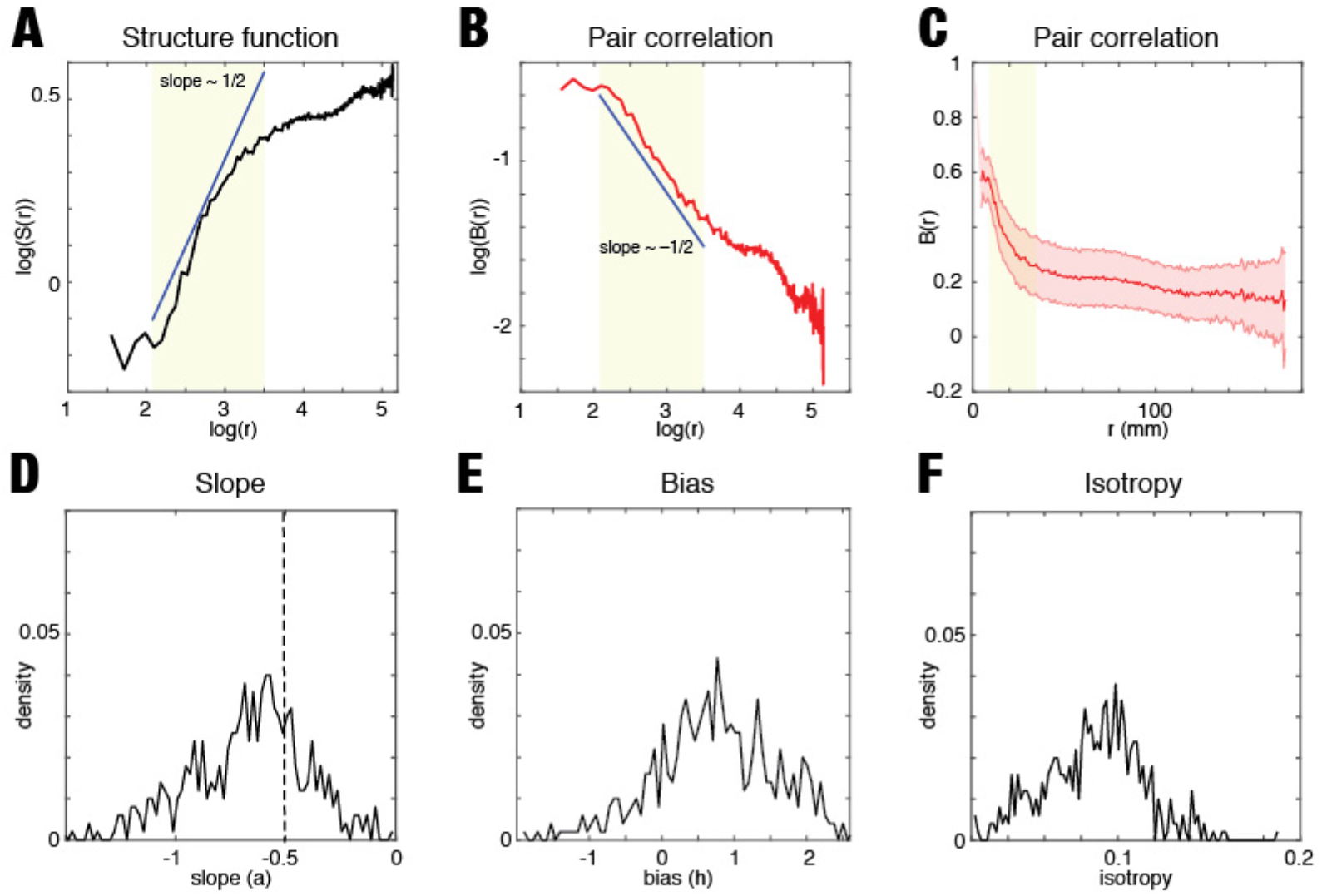
Power laws of the functional core. The figure demonstrates the presence of power law and homogenous isotropy in the empirical human neuroimaging data from 1003 participants. **A)** Spatial power scaling law of the structure function S(r) as a function of log(r) for the correlation function. **B)** Same spatial power scaling law for the correlation B(r) as a function of log(r). **C)** The correlation function B(r) as a function of the distance r, but showing the dispersion across regions. **D)** The unimodal density distribution (across participants and node locations) of the slope parameter a. **E)** Similar unimodal density distribution of the bias parameter h. **F)** Unimodal density distribution of the mean (across r in the inertial subrange) of the standard deviation of B(r). These distributions are suggestive, but not proof, of turbulence and of a functional core of homogeneous isotropic function.

The functional correlation between two nodes, B(r), is computed as a function of the distance between those nodes (averaged across nodes and time), i.e. using homogenous isotropy (see **Figure 7B**). Again, we observe a power law, here with a negative exponent of approximately −1/2 in the same inertial subrange in the functional core. **Figure 7C** shows as B(r) as function of the distance (in a normal coordinate system) but averaged across time and showing the dispersion across nodes, i.e. all pairs across all participants. The unimodal distribution with a single peak suggests homogeneity across nodes in the functional core.

In fact, the homogeneity can be quantified. For each node location 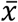 for each participant we compute the fitting of the power scaling law in the same inertial subrange, i.e. log(*B*(*r*)) = *alog*(*r*) + *h*. Here, the parameters *a* (slope) and *h* (bias) describe the power scaling law for each participant and each location. **Figure 7D** shows the density distribution (across participants and node locations) of the slope parameter α, while **Figure 7E** shows this for the bias parameter h. Both distributions are unimodal, which is suggestive of a core of homogeneity of the correlation function B(r) across node location. Furthermore, **Figure 7F** shows the density distribution (across participants and node locations) of the mean (across r in the inertial subrange) of the standard deviation of B(r) (labelled isotropy in the figure). This distribution reflects also a unimodal distribution suggestive of isotropy, given that the variability across directions (standard deviation of B(r)) is consistent with an isotropic peak.

Furthermore, we also checked for power laws in the dynamics of the Hopf whole-brain model. **Figure S3** shows the goodness of fit of power law for the pair correlation function B(r) in the inertial subrange. This is not particularly sensitive to finding optimal turbulence as shown by the red line, representing the goodness of fit, which only goes below p<0.05 for G>0.65. Still, this is evidence for the existence of power law and is consistent with the findings of high values of amplitude turbulence in the same range.

Additionally, in the context of the Exponential Distance Rule, we were interested in estimating the exponential decay, λ, in a complementary way to estimating this from dMRI tractography which yields *λ* = 0.18 mm^−1^. This is consistent with research showing that smaller brain species such as non-human primates and rodents have higher exponential decays (Horvát et al., 2016), with values of *λ* = 0.19 mm^−1^ for non-human primates (Ercsey-Ravasz et al., 2013) and *λ* = 0.78 mm^−1^ for rodents (Horvát et al., 2016). Nevertheless, in order to see if lower values of lambda are feasible in humans, we constructed a Hopf whole-brain model entirely reliant on the Exponential Distance Rule and estimated the possible exponential decays (see Methods). Thus, this model has two free parameters *G* and *λ*, which can be systematically varied to study the root squared error between the empirical and simulated *B*(*r*) in the inertial subrange. **Figure S4A** shows that the model fitting includes the previously empirically estimated *λ* = 0.18 mm^−1^, based on the dMRI connectivity. Importantly, as can be seen from the plot, many exponential decays are possible, including smaller ones as suggested by the existing empirical data in other species.

Similarly, **Figure S4B** shows there is a combination of the two parameters that provides the optimal balance between segregation and integration, computed as the product of the segregation and integration for the model (*see Methods*). **Figure S4C** shows the fit between empirical and simulated B(r) for the inertial subrange, with empirical data shown by the red line and standard deviation, while the data from the Hopf whole-brain model is shown by the blue line and standard deviation. This causally demonstrates that the human brain contains a homogeneous isotropic functional core, which observes spatial power scaling behaviour in the inertial subrange, generated by the Exponential Distance Rule.

Summing up, the results show that the functional core of the human brain exhibits a power law and isotropic homogeneity; both are characteristics of turbulence, and, importantly, this could reflect the presence of an information cascade.

## Discussion

Overall, we used a large, high-quality state-of-art dataset of 1003 HCP participants to demonstrate that human brain dynamics exhibit turbulence as formalised by Kuramoto in his studies of oscillators. Deepening our understanding the causal mechanistic root of this, we built a whole-brain model with coupled oscillators and demonstrated that the best fit of the Hopf whole-brain model to the empirical data corresponds to a region of maximally developed amplitude turbulence. Furthermore, the Hopf whole-brain model shows the economy of anatomy by using the Exponential Distance Rule of anatomical connections as a cost-of-wiring principle. Remarkably, the optimum of turbulence in the model also corresponded to maximal information capability, i.e. sensitivity to the processing of external stimulations, which suggests that turbulence is crucial for information processing.

Probing further the link to information processing in resting state dynamics, we investigated the brain dynamics during seven cognitive tasks in the same participants. We found that the tasks share a turbulent functional core with the resting state, and that the long-distance correlations show task-specific increases in specific higher-order brain regions outside this functional core. We were interested to further establish a link to turbulence in fluid dynamics, and were able to demonstrate a consistent power law for functional brain correlations in a broad spatial range in the functional core suggestive of a cascade of information processing. Overall, our results reveal a way of analysing and modelling whole-brain dynamics establishes turbulence in the dynamic intrinsic backbone of the brain.

### Evidence of turbulence in empirical brain dynamics

The study of turbulence was pioneered by the phenomenological theory of Kolmogorov, based on the concept of structure functions (Kolmogorov, 1941a, b). Equally, Kuramoto were able to formalise a framework for turbulence with a central role for oscillators that is able to model turbulence in fluid dynamics (Kuramoto, 1984). This inspired us to combine Kolmogorov’s structure functions with Kuramoto’s local order parameter to demonstrate turbulence in human brain dynamics. More specifically, we obtained significant results when computing amplitude turbulence is defined as the standard deviation of the modulus of the local Kuramoto order parameter across time and space for the empirical brain resting data compared to applying this to carefully constructed surrogate data. We visualised the change over time and space of amplitude turbulence on a flatmap rendering of the individual empirical brain data. This closely resembled the typical turbulence found in fluid dynamics and oscillators (Kawamura et al., 2007).

Please note that our demonstration of a turbulent dynamic backbone in empirical brain dynamics is entirely compatible with the rich literature on structured temporal patterning in brain data. This can be appreciated by considering two complementary perspectives on brain function, namely *computational* and *dynamical*. The former establishes a relationship between behaviour and concomitant brain activity, while the latter focuses on the information flow across space-time in order to integrate the processing segregated in different neuronal modules. In other words, the dynamical framework provides a description of the communication between nodes.

Our results described are compatible with an account of structured patterns of computation embedded in an intrinsic backbone regulating the windows of opportunity facilitating the communication necessary for integration. Take as an example how metastable dynamics are not only possible but necessary for implementing computation - and for integrating the corresponding structured spatiotemporal patterns (Roberts et al., 2019; Tognoli and Kelso, 2014). The turbulence demonstrated here uses Kuramoto’s framework and generalises previous work showing metastability in whole brain dynamics (Cabral et al., 2014; Deco et al., 2017c), to show that turbulence results in more, rather than less, structure in the brain.

### Modelling the origin of turbulence

Moving beyond correlation, we built a causal mechanistic Hopf whole-brain model using the exponential distance rule for the anatomical structural connectivity to demonstrate the emergence of turbulence. The whole-brain Hopf model of coupled oscillators was able to produce an excellent fit to the empirical data. The results showed maximal *amplitude turbulence* at the dynamical working point of the whole-brain model. Even more, at this working point, we rendered the spatiotemporal evolution of amplitude turbulence on a flatmap of the cortex, which showed a remarkable similarity to the renderings of the empirical data. Importantly, the renderings of other non-optimal working points of the model look rather different as reflected in the Kuramoto amplitude turbulence definition, eg when using very weak connectivity which results in very weak synchronisation and dissolving the vortex structure in the spatiotemporal evolution of patterns (see **Figure 4C**, G=0).

Furthermore, the results suggest that turbulence could play a crucial role in brain information processing, given that we also found maximal information capability of the Hopf whole-brain model for capturing how different external stimulations are encoded in the dynamics. Importantly, we also found an optimal balance between segregation and integration at this working point of the model. Taken together the findings clearly demonstrate that the human brain is turbulent and this helps to facilitate optimal information processing across scales, suggestive of an information cascade.

Please note that an important caveat to using a causal modelling framework is provided by the seminal work of Judea Pearl (Pearl, 2009). In his book “Causality”, he shows that any framework of causal inference is based on inferring causal structures that are equivalent in terms of the probability distributions they generate; that is, they are indistinguishable from observational data, and could only be distinguished by manipulating the whole system. Nevertheless, our modelling framework is perfectly suited for our stated aim of determining the origin of turbulence using a causal modelling framework.

### Task-specific higher-order regions

In order to study the link between turbulence and information processing, we used the same framework to contrast the brain dynamics in seven cognitive tasks with resting state in the same participants. We found that the task-specific functional differences were mainly found in regions in the long-distance subrange of correlations, while the turbulent functional core in the inertial subrange was largely unaffected. This reveals the existence of a turbulent functional core which could be essential for basic brain function and is reflecting the underlying economy of anatomy that keeps the human brain cost effective.

The turbulent functional core is consistent with the discovery of the default mode network (Raichle et al., 2001). This follows the fact that the brain is clearly hierarchical in its structure from single units to the larger circuits (Bullmore and Sporns, 2012; Felleman and Van Essen, 1991; Hagmann et al., 2008; Markov et al., 2014; Mesulam, 1998; van den Heuvel and Sporns, 2011; Zamora-Lopez et al., 2010). In particular, research by Margulies and colleagues (Margulies et al., 2016) have used neuroimaging to extend Mesulam’s seminal proposal that brain processing is shaped by a hierarchy of distinct unimodal areas to integrative transmodal areas (Mesulam, 1998). More recently, we have added to this literature by identifying the ‘global workspace’ of brain regions at the top of the hierarchy (Deco et al., 2020).

Beyond the functional core, the regions that we have identified could promote higher brain function through the breaking of the homogeneity and isotropy of the functional core organisation mainly due to the brain networks found in long-distance subrange, which are the functional homologues driven by the anatomical exceptions to Exponential Distance Rule.

### Finding a power law in the functional core

It is well-known that human brain activity reflects the underlying brain anatomy (Deco et al., 2017c), and that this shaping of function by anatomical connectivity gets even stronger in brain states such as deep sleep (Tagliazucchi et al., 2016) and anaesthesia (Barttfeld et al., 2015). Over the last decades, a large body of convincing research has identified how precisely the underlying anatomical connectivity is responsible for the emergence of the fundamental resting state networks that give rise to the low dimensional manifold of the functional organisation shaped by the human brain (Damoiseaux et al., 2006).

The important result presented here, namely the discovery of a turbulent functional core suggests an even simpler underlying backbone for information processing that can create the necessary efficient information cascade. Supporting this proposal, we were able to show the existence of a power law in the common functional core in the empirical data of both resting state and seven tasks. Taken together, this provides consistent evidence in support of our main findings of turbulence in human brain dynamics as demonstrated using Kuramoto’s oscillator framework in the empirical data and in the Hopf whole-brain models of the data.

The discovery of the turbulent functional core opens up for an elegant proposal, namely that anatomical connectivity of the brain can be described by a structural core following a simple, homogeneous isotropic rule, namely the Exponential Distance Rule that provides the economy of anatomy as a cost-of-wiring principle (Markov et al., 2013).

### Future perspectives

Using state-of-the-art neuroimaging data from over 1000 people, we demonstrate turbulence in human brain activity, in a tour-de-force technical analysis combining empirical methods and wholebrain modelling adapting established methods from the fields of fluid dynamics and oscillators.

This result significantly expands on previous research aiming at relating spatiotemporal chaos to brain activity (Babloyantz and Lourenco, 1994; Breakspear, 2017; Freeman, 2000; Honey et al., 2007; van Vreeswijk and Sompolinsky, 1996). Careful mathematical research has suggested that a main difference between spatiotemporal chaos and turbulence is that the latter is primarily needed for the propagation of disturbances and the transmittal of information from one spatial point to the other (Cross and Hohenberg, 1993; Oono and Yeung, 1987). This could offer a tentative answer not only to Heisenberg’s general question of “Why turbulence” but also to the more specific question of why turbulence in the brain. The purpose of turbulence in the brain must be closely linked to catalyse fast and efficient information processing.

The results presented here from our whole-brain modelling of a very large set of empirical human data confirm that the human brain operates in a turbulent regime showing a maximum of amplitude turbulence and information capability and an optimal balance between integration and segregation.

This finding of turbulence in the human brain is important for its controllability, not only in directing task activity but more generally for characterising brain states in health and disease (Deco et al., 2019; Gu et al., 2017; Tu et al., 2018). As such, the findings will allow for much more sensitive and selective biomarkers of brain states and provide important information on how to control brain disorders and find novel, efficient ways to force homeostatic transitions to a healthy state using external perturbations (Deco et al., 2018; Kringelbach et al., 2020; Kringelbach et al., 2007).

## Acknowledgements

G.D. is supported by the Spanish Research Project (ref. PID2019-105772GB-I00 AEI FEDER EU), funded by the Spanish Ministry of Science, Innovation and Universities (MCIU), State Research Agency (AEI) and European Regional Development Funds (FEDER); HBP SGA3 Human Brain Project Specific Grant Agreement 3 (Grant Agreement No. 945539), funded by the EU H2020 FET Flagship program and SGR Research Support Group support (ref. 2017 SGR 1545), funded by the Catalan Agency for Management of University and Research Grants (AGAUR). MLK is supported by the ERC Consolidator Grant: CAREGIVING (n. 615539), Center for Music in the Brain, funded by the Danish National Research Foundation (DNRF117), and Centre for Eudaimonia and Human Flourishing funded by the Pettit and Carlsberg Foundations.

## Author contributions

G.D. and M.L.K. designed the study, developed the method, performed analyses, wrote and edited the paper.

## STAR Methods

### RESOURCE AVAILABILITY

#### Lead Contact

Further information and requests for resources should be directed to and will be fulfilled by the Lead Contact: Morten L. Kringelbach.

#### Materials Availability

The data set used for this investigation was from an independent publicly available dataset of fMRI data, where we chose a sample of 1003 participants selected from the March 2017 public data release from the Human Connectome Project (HCP). From this large sample we further chose to replicate in the smaller subsample of 100 unrelated participants (54 females, 46 males, mean age = 29.1 +/-3.7 years). This subset of participants provided by HCP ensures that they are not family relatives, and this criterion was important to exclude possible identifiability confounds and the need for family-structure co-variables in the analyses.

#### Data and Code Availability

The HCP dataset is available at https://www.humanconnectome.org/study/hcp-young-adult. The code to run the analysis is available on GitHub (https://github.com/decolab/cr-turbulence).

### Experimental models and subject details

#### Neuroimaging Ethics

The Washington University–University of Minnesota (WU-Minn HCP) Consortium obtained full informed consent from all participants, and research procedures and ethical guidelines were followed in accordance with Washington University institutional review board approval.

#### Neuroimaging Participants

The data set used for this investigation was selected from the March 2017 public data release from the Human Connectome Project (HCP) where we chose a sample of 1003 participants. From this large sample we further chose to replicate in the smaller subsample of 100 unrelated participants (54 females, 46 males, mean age = 29.1+/-3.7 years). This subset of participants provided by HCP ensures that they are not family relatives, and this criterion was important to exclude possible identifiability confounds and the need for family-structure co-variables in the analyses.

#### The HCP task battery of seven tasks

The HCP task battery consists of seven tasks: working memory, motor, gambling, language, social, emotional, relational, which are described in details on the HCP website (Barch et al., 2013). HCP participants performed all tasks in two separate sessions (first session: working memory, gambling and motor; second session: language, social cognition, relational processing and emotion processing).

### METHOD DETAILS

#### Neuroimaging acquisition for fMRI HCP

The 1003 HCP participants were scanned on a 3-T connectome-Skyra scanner (Siemens). We used one resting state fMRI acquisition of approximately 15 minutes acquired on the same day, with eyes open with relaxed fixation on a projected bright cross-hair on a dark background as well as data from the seven tasks. The HCP website (http://www.humanconnectome.org/) provides the full details of participants, the acquisition protocol and preprocessing of the data for both resting state and the seven tasks.

#### Preprocessing and extraction of functional timeseries in fMRI resting data

The preprocessing of the HCP resting state and task datasets is described in full details on the HCP website. Briefly, the data is preprocessed using the HCP pipeline which is using standardized methods using FSL (FMRIB Software Library), FreeSurfer, and the Connectome Workbench software (Glasser et al., 2013; Smith et al., 2013). This preprocessing included correction for spatial and gradient distortions and head motion, intensity normalization and bias field removal, registration to the T1 weighted structural image, transformation to the 2mm Montreal Neurological Institute (MNI) space, and using the FIX artefact removal procedure (Navarro Schroder et al., 2015; Smith et al., 2013). The head motion parameters were regressed out and structured artefacts were removed by ICA+FIX processing (Independent Component Analysis followed by FMRIB’s ICA-based X-noiseifier (Griffanti et al., 2014; Salimi-Khorshidi et al., 2014)). Preprocessed timeseries of all grayordinates are in HCP CIFTI grayordinates standard space and available in the surface-based CIFTI file for each participants for resting state and each of the seven tasks.

We used a custom-made Matlab script using the ft_read_cifti function (Fieldtrip toolbox (Oostenveld et al., 2011)) to extract the average timeseries of all the grayordinates in each region of the Schaefer parcellation, which are defined in the HCP CIFTI grayordinates standard space. Furthermore, the BOLD time series were transformed to phase space by filtering the signals in the range between 0.008-0.08 Hz, where we chose the typical highpass cutoff to filter low-frequency signal drifts (Fox et al., 2005), and the lowpass cutoff to filter the physiological noise, which tends to dominate the higher frequencies (Cordes et al., 2001; Fox et al., 2005). We then applied the Hilbert transforms in order to obtain the phases of the signal for each brain node as a function of the time.

We computed the functional connectivity (FC) as the correlation between the BOLD timeseries in all 1000 regions in the Schaefer Parcellation. We then computed the global brain connectivity (GBC) as the node-level FC, or node strength, characterizing the average FC strength for each region (Demirtas et al., 2019; Yang et al., 2016). Thus, node strength is defined as 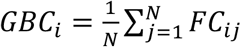.

#### Structural connectivity using dMRI

The Human Connectome Project (HCP) database contains diffusion spectrum and T2-weighted imaging data from 32 participants with the acquisition parameters described in details on the HCP website (Setsompop et al., 2013). The freely available Lead-DBS software package (http://www.lead-dbs.org/) provides the preprocessing which is described in details in Horn and colleagues (Horn et al., 2017) but briefly, the data was processed using a generalized q-sampling imaging algorithm implemented in DSI studio (http://dsi-studio.labsolver.org). Segmentation of the T2-weighted anatomical images produced a white-matter mask and co-registering the images to the b0 image of the diffusion data using SPM12. In each HCP participant, 200,000 fibres were sampled within the white-matter mask. Fibres were transformed into MNI space using Lead-DBS (Horn and Blankenburg, 2016). We used the standardized methods in Lead-DBS to produce the structural connectomes for the Schaefer 1000 parcellation Scheme (Schaefer et al., 2018).

### QUANTIFICATION AND STATISTICAL ANALYSIS

#### Schaefer parcellation

Schaefer and colleagues created a publicly available population atlas of cerebral cortical parcellation based on estimation from a large data set (N = 1489) (Schaefer et al., 2018). They provide parcellations of 400, 600, 800, and 1000 areas available in surface spaces, as well as MNI152 volumetric space. We used here the Schaefer parcellation with 1000 areas and estimated the Euclidean distances from the MNI152 volumetric space and extracted the timeseries from HCP using the HCP surface space version.

#### Analysis using Kolmogorov’ structure function concept

We adapted Kolmogorov’s concept of *structure functions* of a variable *u*, which in turbulence is usually a transversal or longitudinal velocity but here is given by the BOLD signal of the data:

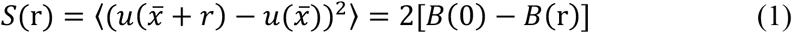

In Equation 1, the basic spatial correlations of two points separated by an Euclidean distance *r*, given by:

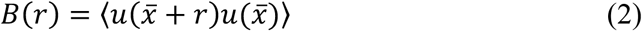

where the symbol 〈 〉 refers to the average across the spatial location 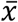 of the nodes and time.

#### Whole-brain model

The link between anatomical structure and functional dynamics, introduced more than a decade ago is at the heart of whole-brain network models (Deco et al., 2013; Deco and Kringelbach, 2014). Typically, the anatomy is represented by the structural connectivity (SC) of an individual or average brain, measured in vivo by diffusion MRI (dMRI) combined with probabilistic tractography. The spatial resolution is in the order of 1-2 mm, but with ultra-high field MRI resolutions 0.4 mm can be reached. The structural connectome denotes the wire-diagram of the connections between cortical regions as ascertained from dMRI tractography. The functional global dynamics result from the mutual interactions of local node dynamics coupled through the underlying empirical anatomical SC matrix. Whole-brain models aim to balance between complexity and realism in order to describe the most important features of the brain *in vivo* (Breakspear, 2017). The most successful whole-brain computational models have taken their lead from statistical physics where it has been shown that macroscopic physical systems obey laws that are independent of their mesoscopic constituents. The emerging collective macroscopic behaviour of brain models has been shown to depend only weakly on individual neuron behaviour. This theoretical framework has been successful in explaining the pattern of inter-regional activity correlation measured with fMRI, so called resting-state-networks. Recent developments have shown that whole-brain models are able to describe not only static FC (averaged over all time points), but also dynamical measurements like the temporal structure of the activity fluctuations, the so-called functional connectivity dynamics (FCD) (Deco et al., 2017c; Hansen et al., 2015).

Here, we use the Hopf model and assume that the underlying anatomy fulfil the *Exponential Distance Rule* derived from exhaustive massive retrograde tract tracing in non-human primates (Ercsey-Ravasz et al., 2013). Mathematically this can expressed as an exponential decay function,

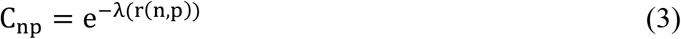

where r(n, p) is the Euclidean distance between the regions n and p, and the decay, λ. Here we estimate λ by fitting the dMRI tractography and obtain λ = 0.18mm^−1^.

The Hopf whole-brain model consists of coupled dynamical units (ROIs or nodes) representing the N cortical brain areas from a given parcellation (Deco et al., 2017c). For the first analysis, we take all 1000 cortical nodes in the Schaefer parcellation. The local dynamics of each brain region is described by the normal form of a supercritical Hopf bifurcation, also known as the Landau-Stuart Oscillator, which is the canonical model for studying the transition from noisy to oscillatory dynamics (Kuznetsov, 1998). Coupled together with the brain network architecture, the complex interactions between Hopf oscillators have been shown to reproduce significant features of brain dynamics observed in electrophysiology (Freyer et al., 2011; Freyer et al., 2012), MEG (Deco et al., 2017b) and fMRI (Deco et al., 2019; Kringelbach et al., 2020).

The dynamics of an uncoupled brain region n is given by the following set of coupled dynamical equations, which describes the normal form of a supercritical Hopf bifurcation in Cartesian coordinates:

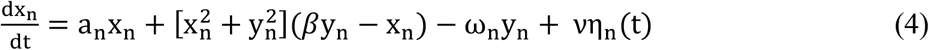

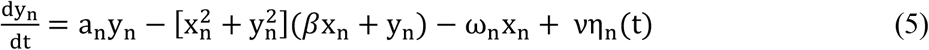

where η_n_(t) is additive Gaussian noise with standard deviation v, and β is the so-called shear factor (where β=0, except in results presented in **Figure S2**, where we systematically explore the influence of this parameter). This normal form has a supercritical bifurcation a_n_=0, so that if a_n_>0, the system engages in a stable limit cycle with frequency *f_n_* = ω_*n*_/2*π*. On the other hand, when a_n_<0, the local dynamics are in a stable fixed point representing a low activity noisy state. Within this model, the intrinsic frequency ω_n_ = ω′_n_ + β, where ω′_n_ is estimated from the empirical data as the peak of the power spectrum. Here, the subindex n denotes the region taken from (1..N), where N is the total number regions.

The whole-brain dynamics was defined by the following set of coupled equations:

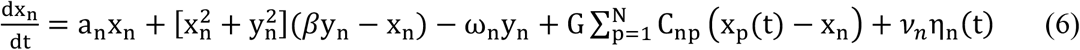

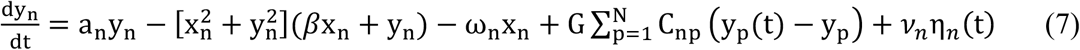

where the noise was fixed v =0.01. The local bifurcation parameters, a_n_ = −0.02, are at the brink of the local bifurcations which is where the best fitting were demonstrated to be achieved. We estimated the intrinsic frequencies from the empirical data, as given by the averaged peak frequency of the narrowband BOLD signals of each brain region. The variable x_n_ emulates the BOLD signal of each region n. To model the whole-brain dynamics we added an additive coupling term representing the input received in region n from every other region p, which is weighted by the corresponding structural connectivity. In this term, G denotes the global coupling weight, scaling equally the total input received in each brain area. All the measures related to whole-brain model were estimated for each global coupling work point, G, running the simulations a 200 times and averaging the results.

#### Measure of amplitude turbulence

We measure amplitude turbulence by first defining the Kuramoto local order parameter and then taking the standard deviation of the modulus across time and space (similar to (Kawamura et al., 2007)). First, we define the amplitude turbulence, *R_n_* (t), as the modulus of the local order parameter for a given brain node as a function of the time:

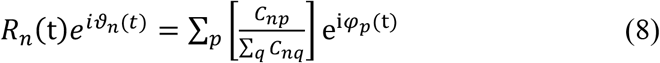

where *φ_p_* (t) are the phases of the BOLD time series and *C_np_* the anatomical exponential distance rule connectivity matrix (see Equation 3). The BOLD fMRI time series were transformed to phase space by first filtering the signals in the range between 0.008-0.08 Hz and using the Hilbert transforms to extract the evolution of the phases of the signal for each brain node over time.

We then measure the amplitude turbulence, D, as the standard deviation across time and space of B:

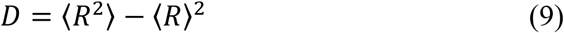

where the brackets 〈 〉 denotes average across space and time. In order to normalise this measure, we shift D with respect to its value when using a global coupling of G=0, which corresponds to a non-coupled system of oscillators, i.e. random.

#### Measure of susceptibility

We define the susceptibility of a whole-brain model as the sensitivity of the brain to the processing of external stimulations. We perturb the Hopf whole-brain model at each G by randomly changing the local bifurcation parameter, a_n_, in the range [-0.02:0]. We estimate the sensitivity of these perturbations on the spatiotemporal dynamics by measuring the modulus of the local Kuramoto order parameter, i.e. 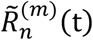 for the perturbed case, and 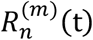 for the unperturbed case. We define susceptibility in the following way:

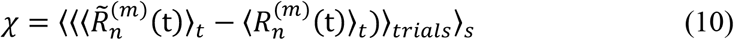

where 〈 〉_*t*_ 〈 〉_*triais*_ and 〈 〉_*s*_ are the mean averages across time, trials and space, respectively.

#### Measure of information capability

Moving beyond susceptibility, we define the *information capability* of the whole-brain model as a measure to capture how different external stimulations are encoded in the dynamics. Specifically, we perturb the model as above, but here the information capability I is defined as the standard deviation across trials of the difference between the perturbed and unperturbed mean of the modulus of the local order parameter across time, averaged over all brain nodes n, i.e.:

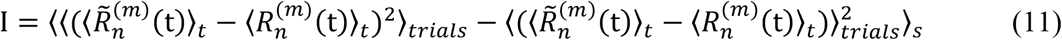

where the averages (〈 〉_*t*_ 〈 〉_*triais*_ and 〈 〉_*s*_) are defined as above.

#### Measure of integration

As a measure of integration we used the mean value of all functional correlation pairs *i* and *j*, i.e.

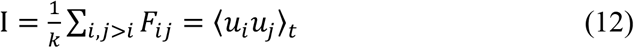

where *k* is the number of upper triangular elements in the functional connectivity matrix *F*, whose elements are defined as the temporal average of the z-scored functional signals *u* between nodes *i* and *j*.

#### Segregation

As a complement of the integration, we used the modularity measure (Rubinov and Sporns, 2011) as a measure of segregation. Following Rubinov and Sporns (2011), modularity is defined as a measure of the goodness with which a network is optimally partitioned into functional subgroups, i.e. a complete subdivision of the network into non-overlapping modules, and supported by densely connected network communities. We consider the modularity of our FC matrix. Our measure of modularity is given by,

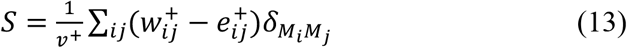

Where the total weight, *v*^+^ = ∑_*ij*_*w_ij_*^+^, is the sum of all positive or negative connection weights (counted twice for each connection), being *w_ij_*^+^ ∈ (0,1] the weighted connection between nodes *i* and *j*. The chance-expected within-module connection weights 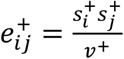, where the strength of node *i*, *s_i_^+^* = ∑_*j*_*w_ij_*^+^, is the sum of positive or negative connection weights of *i*. The *δ_MiMj_ = 1* when *i* and *j* are in the same module and *δ_MiMj_ = 0*, otherwise (Newman, 2006). For a complete description see (Sporns, 2010).

**Supplementary Videos S1-S5. Spatiotemporal evolution of amplitude turbulence in empirical and simulated data.** The videos visualise the change over time and space of the local Kuramoto order parameter, R, reflecting amplitude turbulence in the brain. The videos show the inflated 3D side and midline views as well as flat maps of the hemispheres over the 1200 timepoints of the full resting state session. Video S1 shows the empirical data from a representative single participant, while Videos S2-S5 show the amplitude turbulence for four different values of the global coupling G in the wholebrain model (G=0.8 [optimal], G=0, G=0.4, G=3.0). Related to Figures 2 and 4.

